# NLX-112 is anti-inflammatory, upregulates GDNF and is neuroprotective against MPTP-induced nigrostriatal dopaminergic degeneration in mice

**DOI:** 10.1101/2025.09.25.678507

**Authors:** W. H. Powell, L. E. Annett, R. Depoortere, A. Newman-Tancredi, M. M. Iravani

**Affiliations:** Department of Department of Clinical, Pharmaceutical & Biological Science, Medicine and Life Sciences, College Lane, University of Hertfordshire, AL109AB, UK; Department of Psychology, Sport and Geography, School of Health, Medicine and Life Sciences, College Lane, University of Hertfordshire, AL109AB, UK; Neurolixis SAS, 2 Rue Georges Charpak, 81100, Castres, France

**Keywords:** Neuroprotection, NLX-112, 5-HT_1A_, MPTP, GDNF, open field, thigmotaxis, locomotor activity, mouse model of Parkinson’s disease, microglia, astrocytes

## Abstract

NLX-112 is a potent and selective 5-HT1A agonist that has successfully completed phase 2A clinical trial for treatment of L-DOPA-induced dyskinesia in Parkinson’s disease (PD). We investigated the neuroprotective activity of NLX-112 in a mouse 1-methyl-4-phenyl-1,2,3,6-tetrahydropyridine (MPTP) model of PD. Four groups of mice received either saline (1ml/kg) daily for 15 days, MPTP (21.4 mg/kg for 5 days, preceded and followed by 5 days saline, NLX-112 (1mg/kg/day for 15 days) or combined MPTP + NLX-112. Two weeks following cessation of treatments, NLX-112-treated mice showed increased locomotor activity and reduced anxiety-like behaviour in an open-field test, consistent with sustained effects of 5-HT_1A_ receptor activation. MPTP-treated male mice showed a significant reduction of dopaminergic (i.e., tyrosine hydroxylase immunoreactive; TH-ir) neurones in the substantia nigra (SN) and the striatum by 40 and 55%, respectively. NLX-112 treatment elicited a significant protection against MPTP-induced loss of TH-ir neurones and nerve terminals. MPTP also a markedly increased the levels of GFAP-ir astrocytes and Iba1-ir microglia in the SN, and co-expression of glial-derived neurotrophic factor (GDNF) in the GFAP-ir astrocytes in both the SN and the striatum. However, in MPTP treated mice, NLX-112, markedly reversed microglial expression in the SN, and upregulated GFAP/GDNF co-localisation in both the striatum and the SN. Overall, the present study demonstrates a robust neuroprotective effect of NLX-112 in a mouse model of PD by preventing microgliosis, upregulating GDNF and favouring sustained pro-locomotor activity.

## Introduction

NLX-112 is a first-in-class, highly selective and potent 5-HT_1A_ receptor agonist which counteracts dyskinesia and abnormal involuntary movements in levodopa-primed rodent and primate models of Parkinson’s disease (Iderberg et al., 2015; McCreary et al., 2016; Fisher et al., 2020; Depoortere et al., 2020). Recently, NLX-112 successfully completed a double-blind, placebo-controlled, randomised phase 2A clinical trial showing robust antidyskinetic and antiparkinsonian efficacy (Svennigsson et al., 2025).

There is no known disease modifying treatment for PD so there is a pressing need for discovery of pharmacological neuroprotective agents capable of slowing or preventing the progressive loss of dopaminergic neurons in the substantia-nigra (SN). One potential strategy for neuroprotection is targeting the serotonergic system in the basal ganglia (Di Matteo et al., 2008), and there is strong evidence that drugs which selectively target the serotonin 5-HT_1A_ receptor possess neuroprotective properties. For example, pre-treatment with the 5-HT_1A_ receptor agonist, 8-OH-DPAT, preserves axonal density and protects against N-methyl-D-aspartate (NMDA)-induced cell death in the rat magnocellular nucleus basalis (Oosterink et al., 1998). 8-OH-DPAT also inhibits H_2_O_2_-induced death of rat cortical neurons by suppressing Na^+^ and Ca^2+^ influx, thereby blocking glutamate release (Melena et al., 2000; Lee et al., 2005). Similarly, 8-OH-DPAT almost completely suppresses NMDA-induced caspase-3 activity and DNA fragmentation (Madhavan et al., 2003). The neuroprotective mechanisms of 5-HT_1A_ receptor activation may involve increased phosphorylation of the transcription factor STAT-3, as shown in Neuro2A cells transfected with 5-HT_1A_ receptors where enhanced neurite outgrowth was observed (Fricker et al., 2005). Moreover, repinotan (Bayx3702), which is also a 5-HT_1A_ receptor agonist, inhibits apoptosis in chick embryonic neurons via the nerve growth factor (NGF) signaling pathway (Ahlemeyer et al., 1999), and Bezard et al. (2006) demonstrated that the 5-HT_1A_ agonist BAY639044 is neuroprotective in mice and macaques lesioned with 1-methyl-4-phenyl-1,2,3,6-tetrahydropyridine (MPTP). Both 8-OH-DPAT and repinotan block H_2_O_2_-induced cell death in HN2-5 mouse hippocampal neurons through MAPK activation, suppressing caspase 3 (Adayev et al., 2003). Additionally, 8-OH-DPAT, and the 5-HT_1A_ partial agonists, buspirone and ipsapirone, likely induce protective effects through PI-3K and MAPK activation (Druse et al., 2005).

The neuroprotective role of 5-HT_1A_ receptor agonists is closely tied to astroglial activity (Miyazaki et al., 2020). Mice pre-treated with 8-OH-DPAT and exposed to non-lethal sarin doses exhibit a reduced density of astrocytes (i.e., glial acidic fibrillary protein positive; GFAP-ir cells) in the hippocampus, which correlated with lower levels of interleukin 1 beta (IL-1β) in the baso-lateral amygdala (Garrett et al., 2013). Additionally, 8-OH-DPAT shields rat mesencephalic neurons from 6-OHDA toxicity by enhancing metallothionein in striatal astrocytes (Miyazaki et al., 2013), with astrocyte-induced metallothionein secretion directly protecting dopaminergic neurons from rotigotine toxicity (Isooka et al., 2020).

Overall, the neuroprotective properties of 5-HT_1A_ receptor activation are well-validated but there is no potent, selective and efficacious agonist with regulatory approval. NLX-112 (a.k.a. befiradol or F13640), a selective 5-HT_1A_ agonist with nanomolar affinity which acts as a biased agonist, favoring Gαo protein activation (Colpaert et al., 2002; Newman-Tancredi et al., 2017; 2019), is being developed for symptomatic treatment of motor deficits and levodopa-induced dyskinesia in PD. NLX-112’s agonist efficacy at 5-HT_1A_ receptors is comparable to that of serotonin, surpassing that of other 5-HT_1A_ receptor agonists, including 8-OH-DPAT and buspirone (Newman-Tancredi et al., 2017; 2022).

Since NLX-112 exclusively binds to the 5-HT_1A_ receptor subtype, showing no affinity for dopamine (DA) or other monoaminergic, opioid, GABAergic, or 5-HT receptor subtypes (Colpaert et al., 2002), no known effect on monoamine oxidases, and possesses favorable CNS penetrant pharmacokinetic properties, it has a promising profile as a potential neuroprotective agent in PD. The aim of this study is therefore to investigate whether NLX-112 is also neuroprotective in a commonly utilized model of PD, namely the MPTP mouse model.

## Methods

### Drugs

NLX-112 was provided by Neurolixis and prepared daily in vehicle solution containing saline (0.9% NaCl) and 0.25% dimethyl sulphoxide (DMSO) at a working solution of 1 mg/8 mL. 1-methyl-4-phenyl-1,2,3,6-tetrahydropyridine hydrochloride (MPTP) was purchased from Sigma-Aldrich^®^ (UK) and diluted with saline on the day of experiment. Mice from the MPTP treatment groups were administered 21.4mg/kg MPTP•freebase s.c.) from days 6 to 10. The study involved four groups of mice: (A) vehicle-treated (0.25% DMSO) control mice, (B) NLX-112-treated mice which received daily doses (1 mg/kg i.p.) for 15 consecutive days (day 1 to day 15), (C) mice treated with MPTP, and (D) mice treated with both NLX-112 and MPTP. See Table 1 for dosing regimen.

**Table 1:**
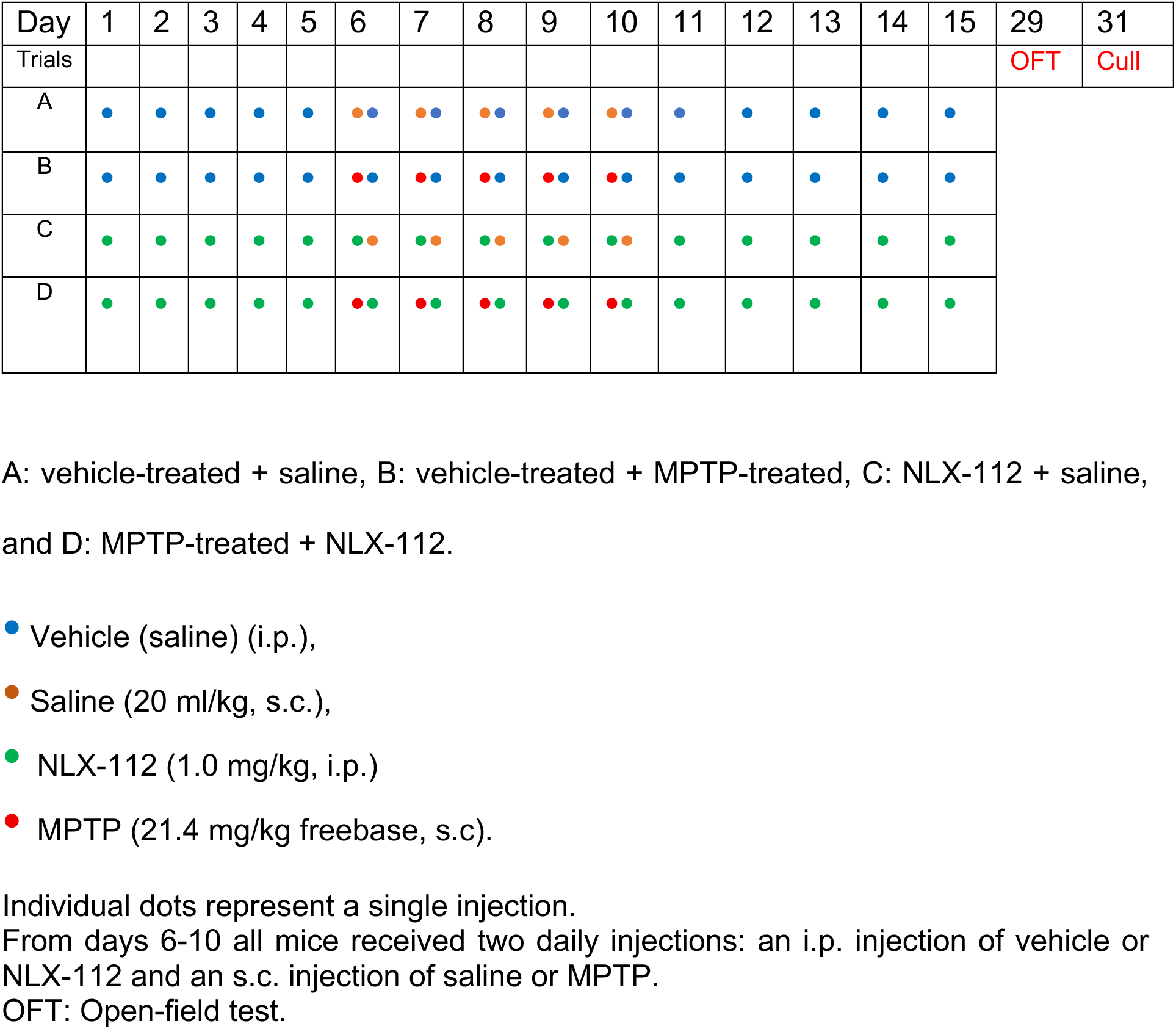
Dosing regimen for treatments:

### Animals

All experiments adhered to the ARRIVE guidelines and were conducted in compliance with the UK Animals (Scientific Procedures) Act 1986 and EU directive 2010/63/EU for animal experiments. The University of Hertfordshire Animal Welfare and Ethical Review Board (AWERB) approved all procedures and protocols, operating under the UK Home Office approved Project License PD7255B4. Male mice (strain: C57blJ6-OlaHSD, RRID:IMSR_ENV:HSD-057), aged 8-10 weeks (19-21g) were purchased from Envigo Laboratories (now Inotiv, UK). This strain of mice is more resilient to the effects of MPTP and exhibits less weight loss during MPTP treatment (Powell et al., 2025, In Press). The sample sizes were chosen based on previous behavioural investigation in mice (Powell et al., 2022, Powell et al., 2025 In Press). Mice were housed in groups of 3 to 4 in single use mouse plastic disposable cages (SUMC, floor area 501 cm^2^; Tecniplast, UK). Access to food and water was ad-libitum, room temperature was maintained at 22°C ± 2°C and light cycle was automated at 12 h: 12 h ((light)/dark cycle). All experiments were carried out under an ambient light intensity of 100 Lux between 09:00 h and 14:00 h.

### NLX-112 and MPTP treatments

Forty-eight adult male C57blJ6-OlaHSD mice were split into 4 groups of equal size: A: vehicle-treated + saline, B: vehicle-treated + MPTP-treated, C: NLX-112 + saline, and D: MPTP-treated + NLX-112. See Table 1 for details.

Because of the risk of contamination, MPTP-treated mice had to be separated from non-MPTP-treated animals. Therefore, it was not possible for effective blinding to take place. However, for assessment of behaviour, animals were arbitrarily assigned blocks in which each block contained either A, B, C and D.

### Video tracking and Open Field test

The open field assessment was performed 14 days after any saline, NLX-112 or MPTP treatment, i.e., just before the mice were euthanised. All open field behavioural assessments were carried out using a Basler infrared camera (Basler AG, Germany), infrared light boxes and analysed with the Noldus EthovisionXT (RRID:SCR_000441) tracking software (Tracksys, Nottingham, UK) according to previously published methods (Powell et al., 2022).

Locomotor activity of mice in an open field test (OFT) was determined by the extent of exploratory movements in a 40 x 40 cm arena. In a novel environment, mice display thigmotaxis, a tendency to remain close to the walls of an arena. To determine the extent of thigmotaxis, a 30 x 30 cm central zone in the centre of the arena was digitally created with the Ethovision-XT software, leaving a 5 cm wide channel around the perimeter of the arena. The following parameters were measured electronically using the Ethovision-XT software: Total distance travelled (cm); stop-start frequency, mean velocity (cm/s), frequency of acceleration in the centre of the arena, entries into the centre arena from the edge, the duration moving into the centre, and the time spent in the centre away from the walls of the arena. Mice were placed in the arena for a total duration of 5 min.

### Immunohistochemistry

Mice were culled by exposure to CO_2_ according to NIH guidelines (CO_2_ flow rate: 5-6 litres/min) in a Vet-Tech medium red chamber (Vet-Tech, UK), transcardially perfused with ice cold phosphate buffered saline (PBS), decapitated, their brains fixed in formalin (10% buffered) and prepared accordingly for immunohistochemical (IHC) analyses. Brains were sectioned coronally at 30 µm thickness and analysed for avidin-biotin visible light IHC for detection of nigral tyrosine hydroxylase (TH)-immunoreactive (-ir) neurons and stereology. Immunofluorescence (IF) immunohistochemistry was carried to detect TH-ir axon terminals in the striatum, astrocytes and microglia using sheep tyrosine-hydroxylase (TH), 1:500 (Thermo Fisher Scientific Cat# PA1-4679, RRID:AB_561880), chicken glial fibrillary associated protein (GFAP) 1:1000 (Thermo Fisher Scientific Cat# PA1-10004, RRID:AB_1074620) and rabbit ionised calcium-binding adapter molecule 1 (Iba1) 1:500 (Abcam Cat# ab178847, RRID:AB_2832244) respectively. We also examined the possibility of glial-derived neurotrophic factor (GDNF) as we have observed previously to be co-localised with astrocytes (Iravani et al., 2012; 2014) using rabbit polyclonal GDNF antibody (Thermo Fisher Scientific Cat# PA5-89957, RRID:AB_2805864). Free floating sections were washed in PBS (1X), triton X-100 (0.1%), non-specific binding was inhibited with non-animal protein blocker (Vector®) and incubated overnight in primary antibody at 4°C. The following day sections were washed and incubated in secondary antibody for one hour at room temp (20-22° C). The relevant secondary antibodies were tagged with AlexaFluor 594 (donkey anti-sheep Thermo Fisher Scientific Cat# A-11016, RRID:AB_2534083, (goat anti-rabbit Thermo Fisher Scientific Cat# A-11012, RRID:AB_2534079) and AlexaFluor 488 (goat anti-chicken Thermo Fisher Scientific Cat# A-11039, RRID:AB_2534096). For avindin-biotin reactions, 3,3-diaminobenzidine (DAB) was used as a chromagen (DAB kit from Vector). Tissue was mounted on slides and images were taken on a Zeiss Axiophot light microscope and photographed with a Zeiss AxioCam SE.

Cell counting in the substantia nigra was performed by either counting every cell that was TH-ir at the levels of the third cranial nerve (Bukhatwa et al., 2009) or the total number of TH-ir neurons were estimated by stereological quantification (Ip et al., 2017). From within the boundaries of the nigrostriatal bundle (∼Bregma −2.50) and the caudal portion of the SNc (∼Bregma −3.90), five TH-ir sections, 300 µm apart (every 10^th^ section) were observed and cells were digitally identified and counted using ImageJ. The following calculation was used to achieve the final estimate of TH-ir neurons.

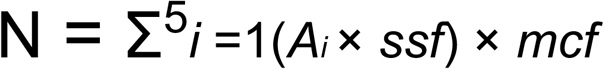

The estimated number of cells (N) has been derived where Σ is the sum of sections*(i)* ranging from 1 to 5 (Σ^5^*i)*. *ssf* (=10) is the sampling fraction which is one section analysed every 10 sections. *mcf* is the missing cell fraction of uncounted cells, cells lay on their side or layered on top of each other through the Z axis = 1.33.

### Optical Density Measurement

To measure the extent of staining TH-ir staining in the striatum, striatal digital images taken at x2 magnification were converted into 8-bit grayscale images and compared against a grayscale stepladder from which a calibration curve was obtained. To assess the extent of nerve terminal loss in the striatum, integrated optical density of striatal sections (3–4 striatal sections per animal) were processed for fluorescence TH-ir. To determine the extent of striatal TH-ir following MPTP or vehicle, the extent of TH-immunostaining was assessed from areas within the striatum using the entire visual field at x20 magnification at 5 to 8 sites. Variations in staining intensity within the dorsal caudate putamen subregions (Iravani et al., 2006) were compared in different animal groups using Image J image analysis software.

### Data Analysis and Statistics

For analysis of each parameter of the OFT, i.e., total distance travelled, frequency of stop-starts, velocity acceleration, entries into the centre and duration of movement in the centre of the arena and for analysis of quantitative immunohistochemical data, we used a mixed model, which allows handling of missing values. When a statistically significant difference was observed in the analysis, data were further analysed using Tukey’s *post-hoc* test. Differences between group means were considered statistically significant when p<0.05. All datasets were tested for normality of distribution using QQ-plots using D’Agostino-Pearson omnibus (K2) test. Normality of distribution was passed when alpha=0.05 or was larger. Statistical analyses were performed using the GraphPad Prism v10.4 (RRID:SCR_002798).

## Results

### Behaviour in the open field

The distance travelled by mice that had received NLX-112+MPTP was more than double that travelled by saline-treated mice [F(4, 50) = 15.76, P< 0.0001, Figure 2A)]. Similarly, the frequency of stop/starts [F(4,46)=18, P<0.0001, Figure 2B], velocity [F(4, 20)=10.83 P<0.0001, Figure 2C], acceleration in the centre of arena [F(4, 51)=11.59, P<0.0001, Figure 2C], entries into centre arena [F(4, 51)=11.53, P<0.0001, Figure 2D] and duration of movement in the centre [F(4, 40)=13.41, P<0.0001, Figure 2F] were all significantly affected by treatments. MPTP treatment significantly slowed vehicle (saline) and NLX-112-treated mice. However, in un-lesioned and MPTP lesioned animals NLX-112-treatment significantly increased distance travelled and stop/start (Figure 2 A,B). Following MPTP, acceleration in the centre of arena, entries into the centre of the arena and duration of movement in the centre of the arena were significantly increased by 1 mg/kg NLX-112 (Figure 2 D-F).

**Figure 2.**
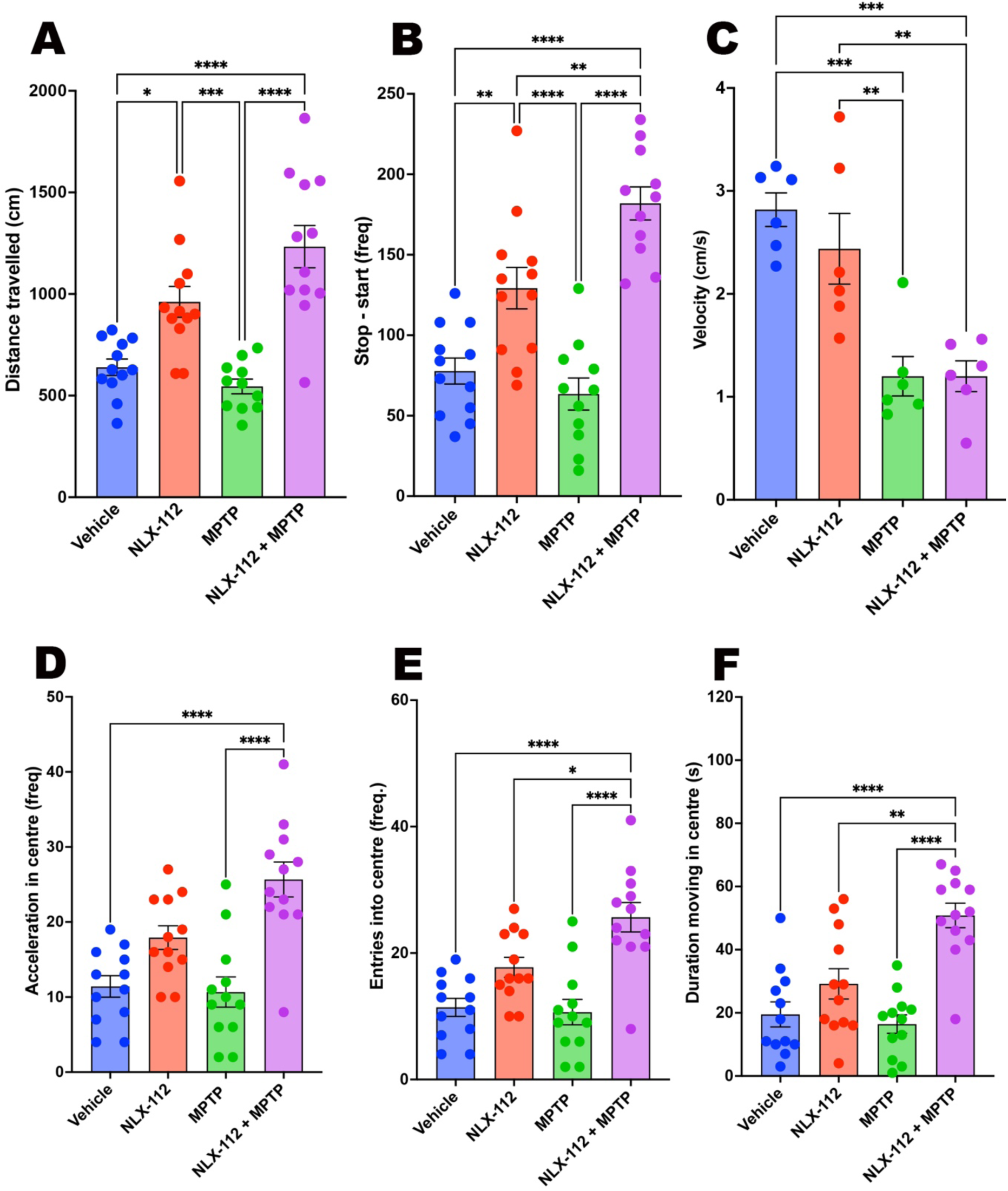
Behaviour over 5 minutes in the open-field test, determined following 14 days of treatment cessation. A) Total distance travelled. B) Frequency of stop / starts, C) Velocity, D) Frequency of episodes of acceleration, E) Frequency of entries into the centre of the arena and F) Time spent in the centre of the arena. . *P<0.05, **P<0.001, ***P<0.0001, Tukey’s *post-hoc* test, following significant One-way ANOVA. N= 10 – 12 per group, Bars are means, error bars are SEMs.

### TH-ir cells and fibres

MPTP treatment led to the death of TH-ir (i.e., dopaminergic) neurons (Figure 3A) and loss TH-ir nerve fibres in the striatum (Figure 3D). Nigral TH-ir neurons were first counted manually at the level of the 3^rd^ cranial nerve. There was an overall statistical difference in number of TH-ir neuron at this level of SNc following different treatments [F(4, 33) = 2.932, P=0.0352]. Treatment with NLX-112 alone had no effect on TH-ir cell number in the SNc or on TH-ir fibre density in the striatum.

**Figure 3.**
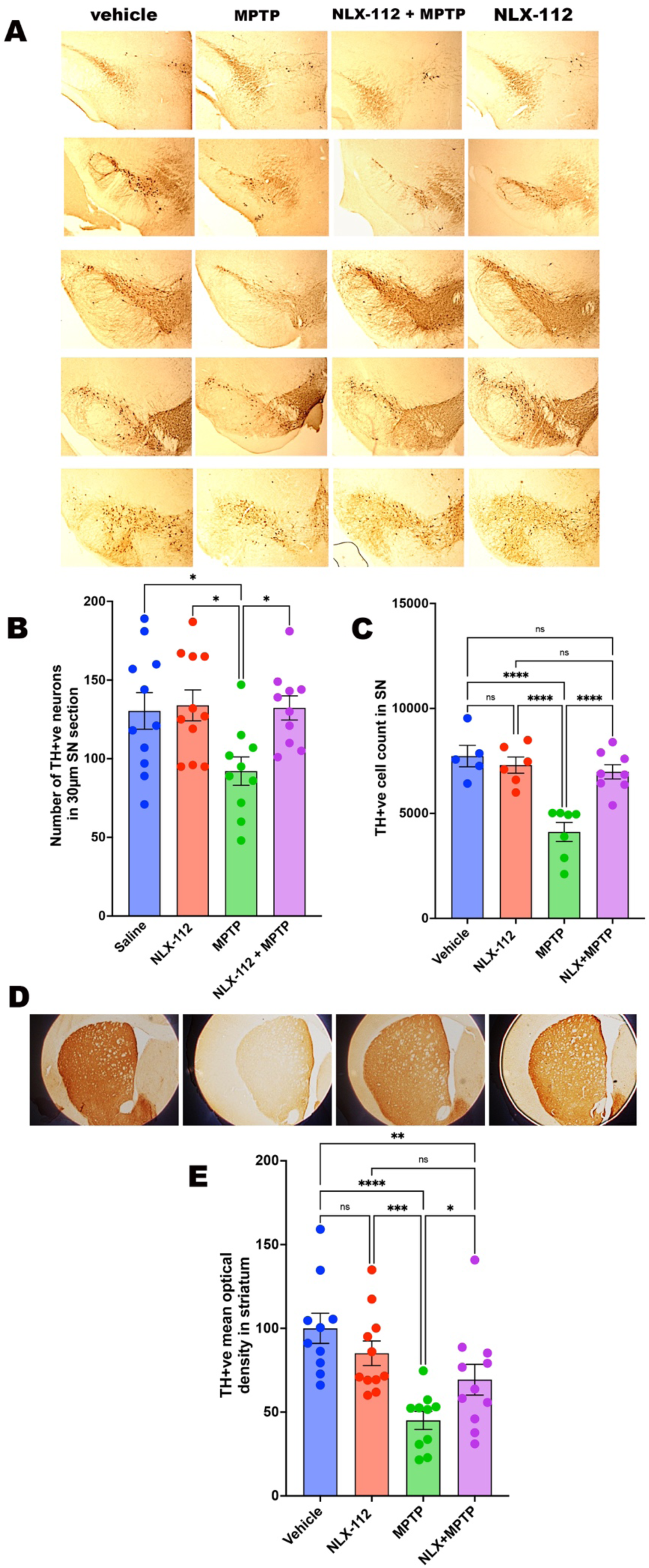
**A)** Representative images of TH-ir stained Substantia nigra (SN) from saline, MPTP, NLX-112 and NLX-112+MPTP mice. Sections were taken from Bregma −2.50 at the nigrostriatal bundle through to Bregma −3.90 at the caudal portion of the SN. **B)** TH-ir neuronal counts at the level of the 3^rd^ cranial nerve. **C)** The number of TH-ir cells stereologically counted in SN from saline, MPTP, NLX-112 and NLX-112+MPTP treated mice and **D)** representative striatal sections from normal, MPTP-treated, contralateral and NLX-112 treated animals. The mean optical density of TH-ir fibres in the CPu is shown in panel **E**. *P<0.05, **P<0.001, ***P<0.0001, Tukey’s *post hoc-*test following significant One-way ANOVA. N= 10 – 12 per group, Bars are means, error bars are SEMs.

NLX-112 1mg/kg/day reversed MPTP induced cell loss of TH-ir neurones (Figure 3B). The total estimate of TH-ir neurons in the SNc following saline, NLX-112, and MPTP+NLX-112 is shown in Figure 3C. MPTP caused 40% loss of these neurons approximately compared to vehicle / saline treatment and saline / NLX-112 groups. Notably, NLX-112 treatment in MPTP group completely protected TH-ir neurons in this brain region [F(3, 22) = 15.67 P<0.0001, Figure 3C]. Protective effects were also observed in the striatum where MPTP caused a 55% loss in TH-ir fibre density, whereas NLX-112 attenuated this loss to 30%, compared to vehicle treatment [F(3, 39) = 8.428, P=0.0002, Figure 3D, E].

### Reactive microgliosis

Immunohistochemical analysis of nigral sections showed presence of numerous Iba1-ir cells in the SN. In the NLX-112 group the number and morphology of these cells did not change. Double immunofluorescence of Iba-1-ir and with TH-ir cells showed that Iba-1-ir cells were much smaller and were interleaved amongst TH-ir neurons. In the MPTP group, loss of TH-ir neurons was co-incidental with an extensive increase in the number of Iba-1-ir microglial cells and change of morphology from fibrillar to a more condensed and occasionally amoeboid shaped. The areas in SNc that were devoid of TH-ir had more activated, condensed and larger Iba-1-ir cells (Figure 4A). In the MPTP + NLX-112 group, the number of activated Iba-1-ir were significantly reduced and the morphology of Iba-1-ir cells were more fibrillar. NLX-112 significantly reduced Iba1-ir in the SN [F(3, 18) = 18.60, P<0.0001, Figure 4 A, B]. Very few Iba-1-ir cells were observed in the striatum.

**Figure 4.**
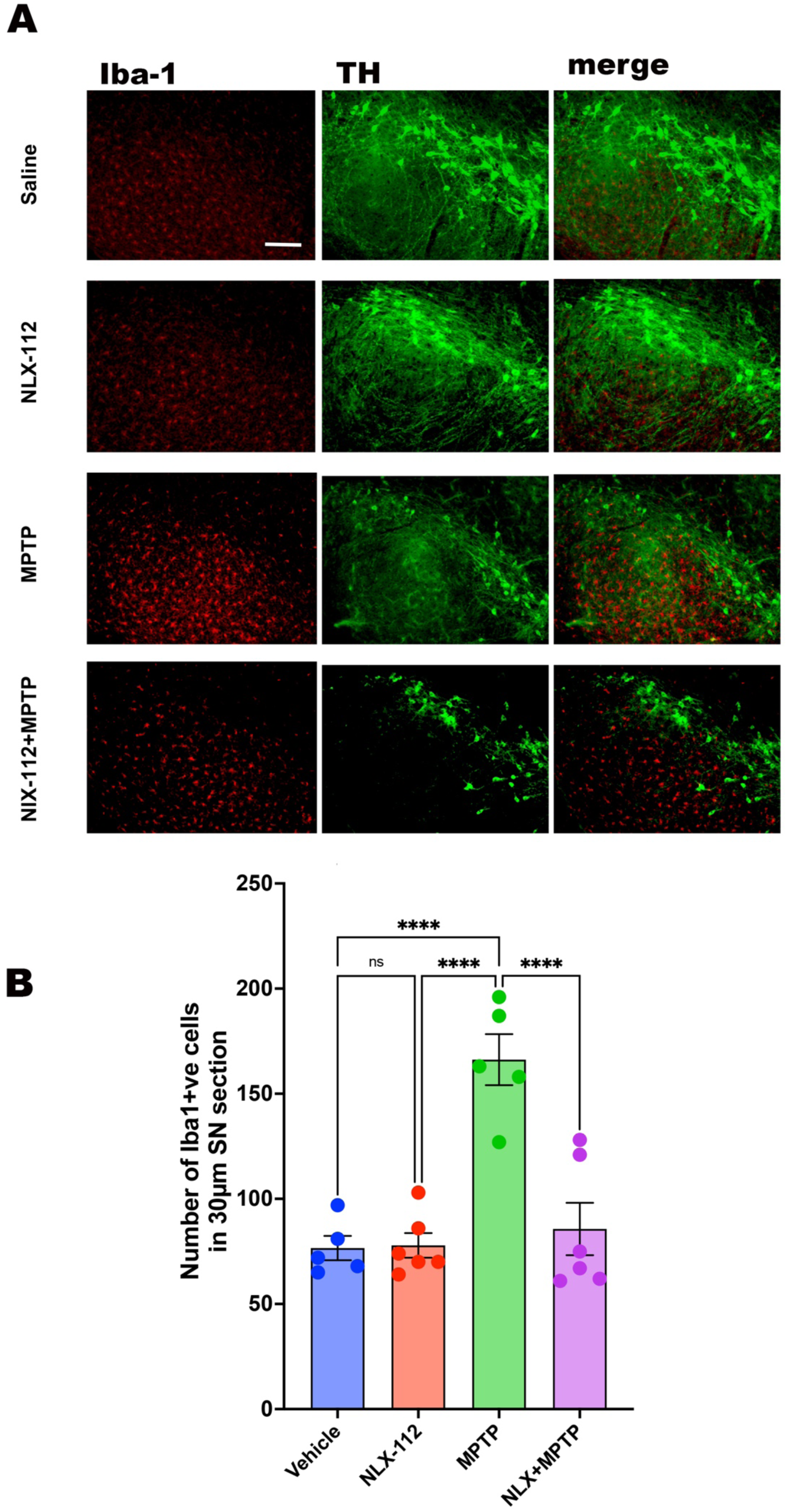
**A)** Reactive micro-gliosis in the SN stained with TH-ir (green) and ionized calcium-binding adapter molecule 1 (Iba1) (red) at X 20 magnification. **B)** Quantification of Iba1-ir cells in the SN. ***P<0.0001, Tukey’s *post hoc-*test following significant One-way ANOVA. N= 5-6 per group, Bars are means, error bars are SEMs. The white scale bar in the upper left photo represents 50 µm.

### Reactive astrocytosis and GFAP-GDNF co-localisation

In the SN of normal mice treated with vehicle (saline) or NLX-112 treated or following MPTP treatment, there was a modest presence of GFAP-ir fibrillar cells. While MPTP did not increase the number of these cells, the morphology of the cells displayed an “activated” state, developing prominent fibrillar end-feet. Double labelling of the nigral sections with glial derived neurotrophic factor (GDNF), showed occasional colocalization of GDNF-ir with GFAP-ir. In MPTP mice that were treated with NLX-112, cells immunoreactive for GDNF appeared to have a distinct astroglial morphology (Figure 5A), more frequently observed [F(3, 35)=9.164 P<0.0001, Figure 5B] and were more frequently associated with GFAP-ir.

**Figure 5.**
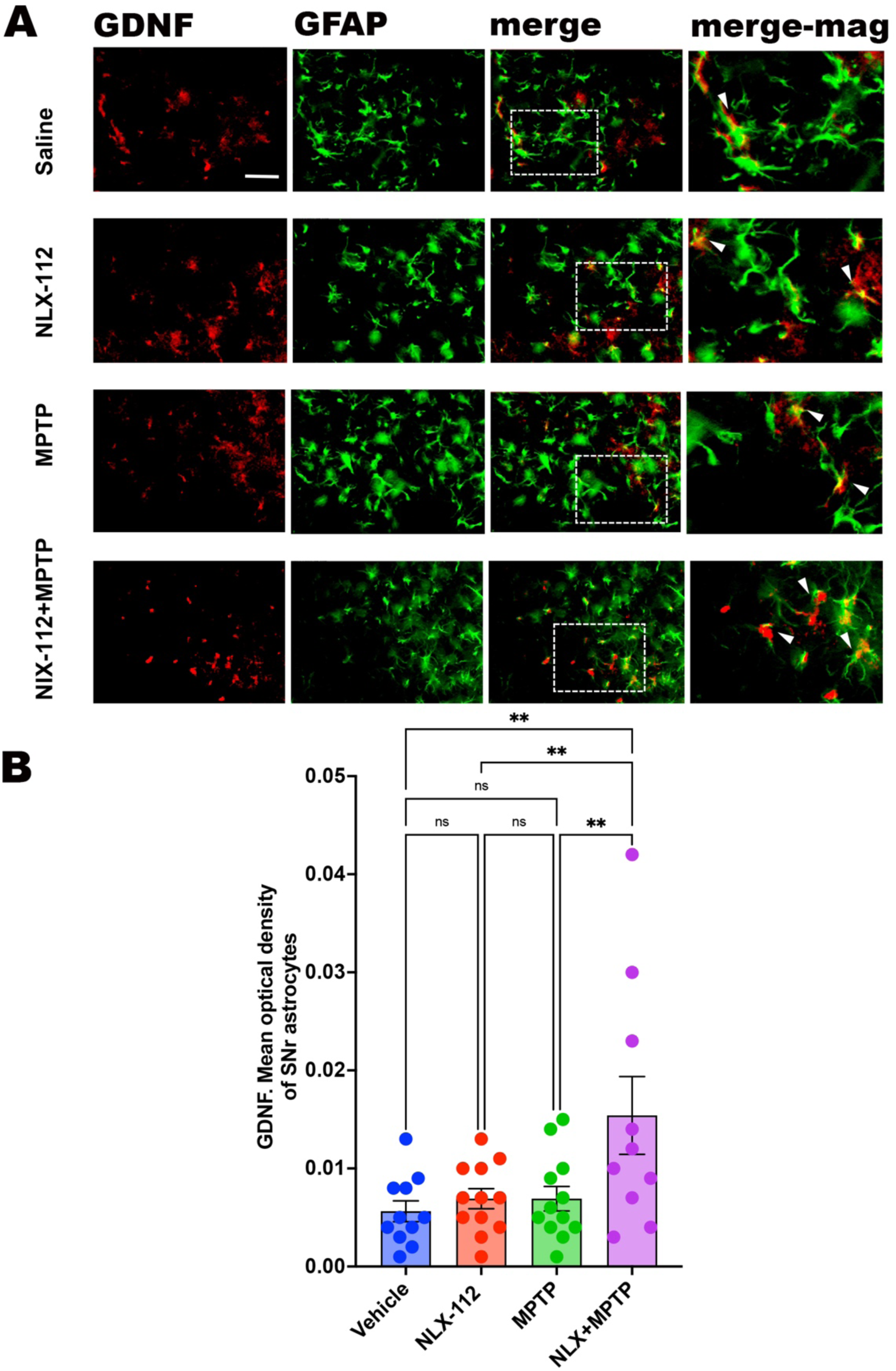
**A)** GFAP and GDNF co-localisation in SN at 40 x magnification. Merge-mag: magnified portion of the merged image (scale bar, 50 µm). **B)** Quantification of co-localisation of GDNF-GFAP in the SNr. One-way ANOVA followed by Tukey’s *post hoc* test. Bars are means, error bars are SEMs. N = 7-12. *P<0.05, **P<0.001, ***P<0.0001, n= 10 - 12, error bars denote SEM. The white scale bar in the upper left photo represents 50 µm.

In the striatum, very few GFAP-ir cells were observed in vehicle (saline) or NLX-112 treated normal mice. GFAP-ir showed extensive co-localisation of GFAP-ir with GDNF-ir (Figure 6A). MPTP treatment markedly increased number of GFAP-ir fibrillar astrocytic cells. In the MPTP-treated animals that received NLX-112, the number of GFAP-ir astrocytic cells decreased [F(3, 35) =11.09 P< 0.0001; Figure 6B]. GDNF-ir immunoreactive cells increased markedly following MPTP and further increase following MPTP =NLX-112 [F(3, 35)=13.44 P<0.0001, Figure 6B]. Co-localisation of GFAP-GDNF in the striatum was increased by 110% following MPTP and was increased even further by 333% in the NLX-112+MPTP group.

**Figure 6.**
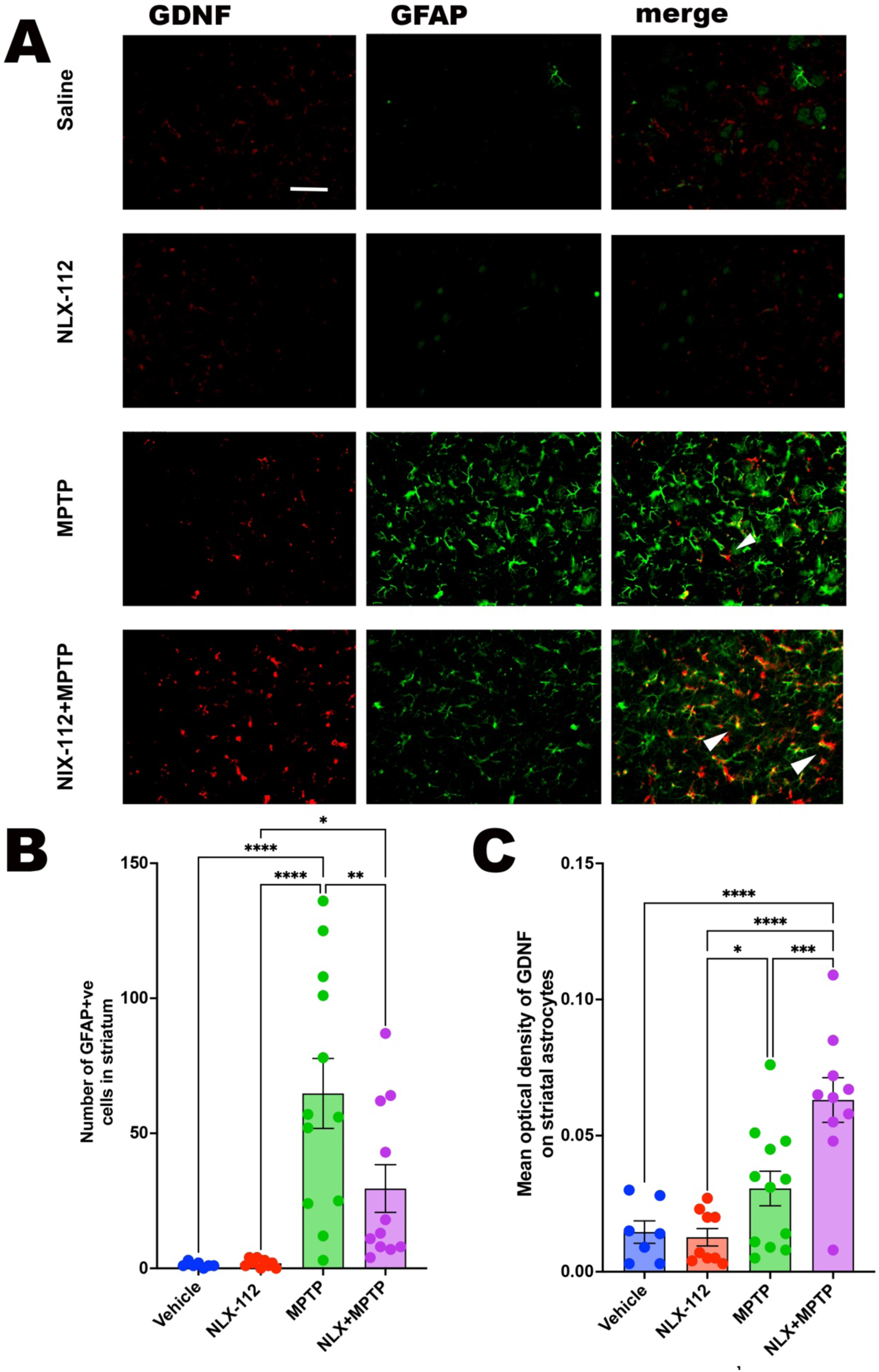
**A)** Representative images of GDNF and GFAP co-localisation in the mid Caudate Putamen of saline, MPTP, NLX-112 and NLX-112+MPTP treated mice, 20X magnification. **B)** Quantification of GFAP immunoreactivity in ventral striatum. **C)** Quantification of GDNF-GFAP co-localisation on striatal astrocytes. One-way ANOVA followed by Fishers LSD. Bars are means, error bars are SEMs. N = 7-12 per group. The white scale bar in the upper left photo represents 50 µm. The white arrows indicate instances of co-localisation.

## Discussion

There is still no treatment that can stop the loss of dopamine-producing brain cells or fully restore their function in Parkinson’s disease. One potential approach involves intraputaminal administration of GDNF which is essential for the growth and survival of these neurons However, clinical trials in humans have had mixed results (Lang et al., 2008; Whone et al., 2019). In one study, low doses of GDNF were intermittently infused and brain scans showed some improvement in dopamine activity, but there was no clear benefit in patients’ symptoms over a 40-week period (Whone et al., 2019). Another study using continuous infusion of GDNF into the brain also failed to show significant improvement compared to a placebo (Lang et al., 2008). In a new approach combing convection-enhanced delivery (CED) — a method designed to more effectively spread treatment throughout the putamen — with gene therapy using AAV2-GDNF (a virus-based system that delivers the GDNF gene) (Heiss et al., 2024; Van Laar et al., 2025). In a recent early-phase trial involving 13 people with advanced Parkinson’s, this treatment showed good safety and was well tolerated (Rocco et al., 2022). Importantly, patients maintained stable motor scores (both on and off medication) over a 45-month follow-up (Heiss et al., 2024).

### NLX-112 is neuroprotective in an MPTP mouse model

The present study shows that the selective 5-HT_1A_ agonist NLX-112: (i) has sustained pro-locomotor behavioural effects consistent with protection of neuronal circuitry involved in motor control, (ii) exerts its neuroprotective efficacy by reducing microgliosis, as MPTP-induced neuroinflammation is markedly reduced, and (iii upregulates endogenous GDNF from astrocytes, and (iv) partially preserves the integrity of DAergic nigrostriatal pathway.

These findings align with an abundance of pre-clinical, as well as clinical research that has sought to harness GDNF as a neuroprotective agent to counter the progressive DA cell loss in PD (Kearns and Gash, 1995; Tomac et al., 1995; Gash et al., 1996; Iravani et al., 2001; Gash et al., 2020; Barker et al., 2020; Manfredsson et al., 2020).

### GDNF

Drinkut et al., (2016) uncovered that the presence and function of the RET receptor was paramount to facilitate the neuroprotective properties of GDNF in the nigrostriatal pathway, by using RET knockout mice that have undergone MPTP challenge. Upon binding to this receptor complex, GDNF initiates signaling pathways that promote neuronal survival, regulate neuronal function, and support the growth and differentiation of specific neuron types, particularly DAergic neurons in the SN (Pascual et al., 2008)

Regarding its relevance to PD, GDNF has garnered meaningful attention due to its potential therapeutic implications. Research from preclinical animal models suggests that GDNF exhibits neuroprotective effects by fostering the survival of DA-producing neurons within the SN (Tomac et al., 1995; Kordower et al., 2000; Sun et al., 2005). Moreover, GDNF injections into the rat SN shield DA neurons in the SN and VTA from 6-OHDA induced cell death (Kearns and Gash, 1995). The protective effect of GDNF was also demonstrated in mice (Tomac et al., 1995) and non-human primates (Gash et al., 1996), where it was also shown to foster protection and regeneration of the DAergic nigrostriatal system upon MPTP lesioning (Iravani et al., 2001), as well as improving motor activity (Costa et al., 2001). Additionally, GDNF likely modulates neuronal function and connectivity, potentially aiding in the restoration of impaired motor function in PD (Kirik et al., 2001; Boger et al., 2006). Furthermore, decreased levels of GDNF in PFC of MPTP treated mice causes synaptic degeneration and is associated with cognitive impairment (Tang et al., 2023). Finally, GDNF is implicated in the genetic predisposition to PD associated with LRRK2 mutations (Jaimon et al., 2025).

However, despite promising preclinical studies demonstrating its neuroprotective and restorative properties, clinical trials exploring GDNF as a direct treatment for PD have yielded mixed results (Gash et al., 2020; Barker et al., 2020; Manfredsson et al., 2020). But, and importantly, these studies were carried out through the administration of exogenous GDNF (Winkler et al., 1996; Georgievska et al., 2002; Sun et al., 2005), GDNF-secreting mesenchymal stem cells (Hoban et al., 2015) or GDNF-transfected macrophages (Biju et al., 2010; Zhao et al., 2014). A fundamental difference that distinguishes the present work (based on targeting 5-HT_1A_ receptors) from the studies mentioned above is that it is based on upregulation of the brain’s own endogenous GDNF, rather than administering GDNF exogenously. This resulted in robust protection of the nigrostriatal pathway and suggests that activation of serotonin 5-HT_1A_ receptors with a selective and efficacious agonist such as NLX-112 could be a promising therapeutic strategy (see below).

### NLX-112 upregulates astrocytic GDNF

The results in the present study revealed that the upregulated GDNF is localised on GFAP-ir cells (i.e., astrocytes). Astrocytes are extremely multifaceted, facilitating the transport of macromolecules from the BBB to resident neurons, and are also one of the principal cell types that secrete GDNF. Thus, knockout of astrocytic SARM1 (a member of the Toll/interleukin receptor family) in mice with experimental autoimmune encephalomyelitis (EAE; a model of Multiple Sclerosis) inhibits neuroinflammation through upregulation of GDNF (Jin et al., 2022). Furthermore, upon ischemic damage, neurons are rescued by high frequency magnetic stimulation through the release of GDNF on astrocytes (Gava-Junior et al., 2023).

In addition, astrocytes are known to express 5-HT_1A_ receptors (Whitaker-Azmitia et al., 1993; Miyazaki et al., 2013; 2016; 2017; Lee et al., 2015; Isooka et al., 2020; Narváez et al., 2020; Kikuoka et al., 2020) and 5-HT_1A_ agonists such as 8-OH-DPAT or ipsapirone have been used to elicit neuroprotection through release of antioxidants such as metallothionine or the cell growth regulating protein, S-100 (Whitaker-Azmitia et al., 1993; Miyazaki et al., 2013; 2016; 2017). The 5-HT_1A_ receptor has also been directly observed by IHC on astrocytes in the gerbil hippocampus but interestingly the 5-HT_1A_ receptor was only shown to be colocalised with astrocytes upon ischemic neuronal damage, and not in normal non-hypoxic astrocytes (Lee et al., 2015). This suggest that astrocytes do not engage 5-HT_1A_ receptors as a basic resting function, but that toxic insult or damage to resident neurons causes 5-HT_1A_ receptors to be expressed and become functionally active. This is supported by the fact that resting astrocytes are known to contain 5-HT_1A_ mRNA (Hirst et al., 1998) but that 5-HT_1A_ agonism using 8-OH-DPAT is unable to inhibit cAMP production, signifying the absence of functional 5-HT_1A_ receptors. These observations suggest that the role of 5-HT_1A_ receptors in astrocytes is dependent on a neurotoxic challenge (e.g. via ischemic damage or using a toxin such as MPTP) which would shift 5-HT_1A_ receptors from a dormant to a functional state which can be activated by agonists such as NLX-112. This is consistent with the fact that astrocytes harbour 5-HT_1A_ mRNA and that NLX-112 elicits a major upregulation of GDNF in mouse nigrostriatal neurons as well as causing a reduction in GFAP-ir astrocytic activation. This is further supported by GDNF release only occurring in response to NLX-112 in the presence of MPTP, but not by either treatment alone (Figure 4).

### NLX-112 neuroprotection: mechanism of action

MPTP crosses the blood brain barrier (BBB), and is taken up by astrocytes and converted to the toxic metabolite MPP^+^ by the monoaminergic enzyme, MAO-B. It is then released into the synaptic cleft where it is taken up into DA neurons via the dopamine transporter (DAT). Once in the cytosol of DA neurons it functionally inhibits mitochondria, resulting in the cessation of ATP production. Loss of mitochondrial function then leads to a build-up of ROS, such as hydrogen peroxidase. The build-up of reactive oxygen species (ROS) and loss of mitochondrial function initiates apoptosis leading to death of the DA neuron. This process triggers astrocytic activation and production of GDNF in response to apoptotic chemokine and cytokine signalling. The present data suggests that NLX-112’s potent agonism at the 5-HT_1A_ receptor leads to the sustained activation of astrocytes, leading to amplified secretion of GDNF. Astrocytic secreted GNDF is then taken up via RET receptors on DA neurons resulting in greater protection of the neuron and its survival against MPP^+^ toxicity. Additionally, the signaling pathway of GDNF/RET helps regulate optimal mitochondrial function and activity through stimulation of complex-I whilst functionally interacting with the NfκB pathway thereby enhancing mitochondrial biogenesis and restoring ATP production (Kramer and Liss et al., 2015)

### NLX-112 facilitates locomotor activity and anxiolytic properties

Importantly, the neuroprotective properties observed *in vitro* were found to translate to behavioral effects *in vivo*. NLX-112 increased locomotor activity and reduced anxiety-like behavior of the mice, effects which were observed 2 weeks after treatment cessation, consistent with long-term neuroprotective and behavioral effects. NLX-112 binds to the 5-HT_1A_ receptor in key brain regions associated with motor function (Newman-Tancredi et al., 2022), particularly throughout the cortical regions and striatum, two areas heavily involved in control of CNS and PNS motor units (Ebbesen and Brecht, 2017). This, paired with the fact that mice were treated with NLX-112 (1mg/kg) for 15 days could have led to changes in synaptic plasticity that protect against MPTP-induced toxicity and facilitate locomotor activity.

In addition to pro-locomotor activity, NLX-112 also increased the amount of time that the mice spent in the centre of the open field arena, consistent with anxiolytic-like properties. In our previous paper (Powell et al., 2022) we observed anxiolytic effects following acute administration of NLX-112 at low doses (0.1, 03 and 1.0mg/kg) in older mice (6 months), in the open-field and elevated plus maze tests. The present study now expands on those findings by showing that repeated administration of NLX-112 has protracted anxiolytic-like activity (increased entries in the centre quadrant) even 2 weeks after treatment cessation. Moreover, this effect is amplified in mice that have received both NLX-112 and MPTP, suggesting that the deleterious effects of MPTP on CNS monoaminergic function cause changes in 5-HT_1A_ receptor sensitivity rendering the mice more responsive to treatment with NLX-112. This hypothesis is supported by rat brain imaging studies in which 5-HT_1A_ receptors were labelled with ^18^F-NLX-112 (a.k.a. ^18^F-F13640) and showed changes in sensitivity following 6-OH-DA lesion and L-DOPA treatment (Vidal et al. 2021). Another brain imaging study using ^18^F-FDG showed that there were broad changes in energy use in hemiparkinsonian (i.e. 6-OH-DA lesioned) dyskinetic rats and these were partially rescued by NLX-112 treatment (Chaib et al. 2023). Overall, these observations suggest that activation of 5-HT_1A_ receptors by NLX-112 elicits neuroprotective and neuroplastogenic effects that safeguard neuronal pathways involved in motor control and provide benefit in models of anxiety/mood.

## Conclusion

This study shows that NLX-112 exhibits neuroprotective properties in a MPTP mouse model of PD. NLX-112’s protective effects are likely mediated through reversal of MPTP-induced inflammation as shown by attenuation of astrogliosis and microgliosis, and, importantly, by upregulation of the neurotrophic factor GDNF in astrocytes. Notably, the neuroprotective effects observed at the cellular level translated to robust and sustained pro-locomotor and anxiolytic-like effects in the open-field behavioural test. These observations suggest that targeting astrocyte-expressed 5-HT_1A_ receptors with a selective and high efficacy agonist could constitute a novel and promising therapeutic strategy for neuroprotection and disease modification in Parkinson’s disease.

## Acknowledgement

This work was supported by European Union Regional Development Fund and Hertfordshire Knowledge Exchange Partnership (HKEP). We thank Mark A. Varney for helpful comments on the manuscript.

## Data Availability

All data generated and analysed is available unrestricted and can be provided upon request.

## Abbreviations

MPTP: 1-methyl-4-phenyl-1,2,3,6-tetrahydropyridine
PD: Parkinson’s disease
TH: tyrosine hydroxylase
GDNF: glial cell-line derived neurotrophic factor
GFAP: glial fibrillary acidic protein
IBA-1: ionized calcium-binding adapter molecule 1

